# Overcoming PARP inhibitor resistance by inducing a homologous recombination repair defective phenotype with ATR, CHK1 and WEE1 inhibitors

**DOI:** 10.1101/2023.07.05.547758

**Authors:** Hannah L Smith, Elaine Willmore, Lisa Prendergast, Nicola J Curtin

## Abstract

**Purpose:** PARP inhibitors (PARPi) are effective in homologous recombination repair (HRR) defective (HRD) cancers. To (re)sensitise HRR proficient (HRP) tumours to PARPi combinations with other drugs are being explored. Our aim was to determine the mechanism underpinning the sensitisation to PARPi by inhibitors of cell cycle checkpoint kinase ATR, CHK1 and WEE1.

**Experimental design:** A panel of HRD and HRP cells (including matched BRCA1 or 2 mutant and corrected pairs) and ovarian cancer ascites cells were used. Rucaparib (PARPi) induced replication stress (RS) and HRR (immunofluorescence microscopy for γH2AX and RAD51 foci, respectively), cell cycle changes (flow cytometry), activation of ATR, CHK1 and WEE1 (Western Blot for pCHK1S345, pCHK1S296 and pCDK1Y15, respectively) and cytotoxicity (colony formation assay) was determined, followed by investigations of the impact on all of these parameters by inhibitors of ATR (VE-821, 1 μM), CHK1 (PF-477736, 50 nM) and WEE1 (MK-1775, 100 nM).

**Results:** Rucaparib induced RS (3 to10-fold), S-phase accumulation (2-fold) and ATR, CHK1 and WEE1 activation (up to 3-fold), and VE-821, PF-477736 and MK-1775 inhibited their targets and abrogated these rucaparib-induced cell cycle changes in HRP and HRD cells. Rucaparib activated HRR in HRP cells only and was (60-1,000x) more cytotoxic to HRD cells. VE-821, PF-477736 and MK-1775 blocked HRR and sensitised HRP but not HRD cells and primary ovarian ascites to rucaparib.

**Conclusions:** Our data indicate that, rather than acting via abrogation of cell cycle checkpoints, ATR, CHK1 and WEE1 inhibitors cause an HRD phenotype and hence synthetic lethality with PARPi.

## Introduction

PARP inhibitors (PARPi) are synthetically lethal in tumours with BRCA mutations or other homologous recombination repair (HRR) defective (HRD) tumours (1, 2), because PARPi cause replication-associated lesions that can only be resolved by HRR (3). Six PARPi are approved for ovarian, breast, pancreatic and castrate-resistant metastatic prostate cancers by the FDA/EMA (rucaparib, olaparib, niraparib and talazoparib) (https://www.fda.gov, accessed 3^rd^ February 2023) and the Chinese NMPA (pamiparib and fuzuloparib) (4, 5). However, despite the efficacy of PARPis in HRD tumours, HRR proficient (HRP) cancers are intrinsically resistant and HRD cancers that are initially responsive can become resistant to PARPi, often due to restoration of HRR (6). Therefore, new ways of improving PARPi therapy are required to overcome both of these issues.

One approach to improving PARPi therapy is by combination with other anticancer agents. However, combinations with cytotoxic drugs and radiotherapy have proved to increase toxicity with insignificant impact on the overall therapeutic response (7, 8). A more promising approach is the combination of PARPi with other molecularly targeted therapies. ATR, CHK1 and WEE1 are key kinases which, in response to replication stress (RS), signal to S and G2/M cell cycle checkpoints to exploit the nearly ubiquitous dysfunction of the G1 checkpoint in cancer, e.g., due to mutations in p53 and pRb (9). As PARPi cause RS, targeting the RS response by combining PARPi with ATR/CHK1/WEE1 inhibition provided a sound rationale for this study. Individually ATR, CHK1 and WEE1 have also each been reported to signal to HRR, with inhibition of each of these kinases impairing HRR (10-12). An early study demonstrated that a prototype ATR inhibitor (ATRi), NU6027, was synergistically cytotoxic with the PARPi, rucaparib, and the major contributor to this synergy was inhibition of HRR (13). Previously we found that the CHK1 inhibitor (CHK1i), PF-477736, sensitised HRP cells to rucaparib, but there was no sensitisation in matched HRD cells, indicating the mechanism of synergy was primarily HRR inhibition (14). Preclinical studies indicate that ATRi, CHK1i and WEE1i enhance PARPi cytotoxicity *in vitro* and *in vivo* (15-23). Several ongoing clinical trials focus on PARPi and ATRi combined (NCT0405269, NCT04267939, NCT04972110, NCT03787680, NCT04149145, NCT03682289, NCT03022409), as well as CHK1i (NCT04030559) and WEE1i (WEE1i) (NCT02576444, NCT03579316). Results to date show promise as results from the CAPRI phase 2 study with olaparib (PARPi) and cerelasertib (ATRi), showed this combination was well tolerated in high-grade serous ovarian carcinoma patients (NCT03462342) (24). Phase 1 trials have shown olaparib is well-tolerated with CHK1i prexasertib and WEE1i adavosertib (MK-1775) (NCT04197713) (25, 26). No preclinical study as yet has directly compared the effects of ATR, CHK1 and WEE1 inhibitors on PARPi cytotoxicity, or the relative contributions of the impact of cell cycle checkpoint signalling and HRR on the sensitisation observed.

The aim of this study therefore was to: i) compare ATR, CHK1 and WEE1 inhibitors to determine which had the greatest impact as single agent, ii) assess which of the inhibitors sensitised cells the most to the PARPi, rucaparib, and iii) investigate the mechanisms underpinning sensitisation. We conducted these investigations in human ovarian cancer cells, where PARPi have the most approvals for clinical use, and HPV positive and negative cervical cancer cells, where PARPi are not currently approved. To determine the relative impact of ATR/CHK1/WEE1 inhibition on HRR vs cell cycle we compared isogenically matched HRP and HRD cells, by virtue of *BRCA1* (UWB cells) or *BRCA2* (V-C8 cells) mutation with the matched corrected cells. This combination of rucaparib with the checkpoint kinase inhibitors was further tested on malignant ovarian ascites cells, characterised for their HRR status, to assess the translational relevance of acquired *in vitro* data. Here, we show that rucaparib activates cell cycle checkpoint kinases and their corresponding inhibitors sensitise HRP but not HRD cell lines or patient cultures to rucaparib. We conclude the primary mechanism of this sensitisation is therefore via the impact on HRR, with ATRi, CHK1i and WEE1i having equivalent effects at inducing an HRD phenotype in HRP cells.

## Methods

### Cell culture

All human cell lines were obtained from American Type Culture Collection (ATCC; Manassas, VA, United States) or the European collection of authenticated cell cultures (ECACC, Salisbury, UK) and were used within 30 passages of purchase or subsequent authentication by: STR profiling (PowerPlex Fusion System, Promega). C33A and SiHa cells were grown in Dulbecco’s Modified Eagle Medium (DMEM;). IGROV-1 cells were cultured in (RPMI)-1640 medium. Paired *BRCA1* mutant UWB1.289 (UWB) and the corrected UWB1.289+ *BRCA1* (UWB+B1) cells generated by transfection with pcDNA3 plasmid containing wild-type *BRCA1* (27) were both grown in 50% RPMI-1640 and 50% Mammary Epithelial Growth Medium (Lonza, Basel, Switzerland). Medium for UWB+ B1 cells was supplemented with 200 μg/ ml G418 S (Merck, Kenilworth, NJ, United States) to maintain the transfection. Chinese hamster lung fibroblasts with mutant *BRCA2* V-C8 and their *BRCA2* corrected sub-line, V-C8.B2 cells, were a gift from Thomas Helleday (28) cultured in DMEM media, with 200 μg/ml G418 S (Merck, Kenilworth, NJ, United States) added to the culture media of the *BRCA2* corrected cells. Cells were maintained in exponential phase at 37 ° C, 5% CO_2_ and 95% humidity. Cell lines were mycoplasma free (MycoAlert, Lonza, Basel, Switzerland).

### Chemicals and reagents

All routine chemicals were obtained from Sigma-Aldrich (St. Louis, MO, USA), unless stated otherwise. The PARPi used in this study, rucaparib, was kindly gifted from Pfizer Global R&D (La Jolla, CA, USA). ATRi VE-821, CHK1i PF-477736 and WEE1i MK-1775 were purchased from Selleckchem (Houston, TX, USA), prior to dilution to stock concentrations of 20 mM or 10 mM in dry DMSO, and storage at -20°C to avoid degradation.

### Colony Formation Assay

Exponentially growing cells were seeded in 6-well plates at densities ranging from 50 to 4000 cells/well, with the aim of having 20-200 countable colonies at the termination of the experiment. After attachment, cells were exposed to 0.5% DMSO (vehicle control) or drugs, alone and in combination in 0.5% DMSO for 24 h then cultured in fresh medium for 10-14 days, allowing colony formation. Colonies were fixed with 3:1 methanol: acetic acid and stained with 0.4% crystal violet then counted by eye. Data were imported into GraphPad Prism 9.0, for calculation of LC_50_ values and statistical analysis.

### Western blotting to determine target inhibition

Cells were seeded at 2-5 x 10^5^ in 10 cm^2^ dishes and, when 70-80% confluent, exposed to media containing drug for 24 hr. Cells were lysed using phosphosafe extraction reagent (Merck, Kenilworth, NJ, United States) with 1:100 protease cocktail inhibitor (Thermo Fisher Scientific, Waltham, MA, United States) at 4 ° C. Cells were scraped, centrifuged and diluted in diH_2_O to 0.5-1 mg protein/ml. XT sample buffer (Bio-Rad Laboratories, Hercules, CA, United States) and XT reducing agent (Bio-Rad Laboratories, Hercules, CA, United States) were added at constants of 25% and 0.5%, respectively. Samples and HiMark pre-stained protein standard (Thermo Fisher Scientific, Waltham, MA, United States) were loaded onto 3-8% Criterion XT tris-acetate gels (Bio-Rad Laboratories, Hercules, CA, United States) which was run at 150V for 1 hr. Separated proteins on the gel were transferred onto nitrocellulose membrane (Amersham) at 100V for 1 hr before exposure to primary antibodies pCHK1S345 (Cell signalling #2348, diluted 1:1000), pCHK1S296 (Cell signalling #90178, diluted 1:1000), pCDK1Y15 (Cell signalling #9111, diluted 1:1000) and vinculin loading control (Cell signalling #4650, diluted 1:1000) in 5% bovine serum albumin (BSA) (Merck, Kenilworth, NJ, United States)/TBS-Tween overnight at 4° C. Following washes, secondary anti-rabbit antibody (Dako #PO447, diluted 1:2000 in 5% milk powder in TBS-Tween) was added before further washes and the addition of Clarity Max ECL (Bio-rad, Hercules, CA, United States) to image membrane sections using the G Box (Syngene, Cambridge, UK).

### Immunofluorescence for assessing RS and HRR function

Replication stress (RS) was measured by γH2AX focus formation and HRR function by RAD51 focus formation. Cells were seeded onto sterile coverslips and left to adhere for 24 h (cell lines) or 48 h (patient ascites) prior to exposure to 10 μM rucaparib single agent for 24 h (cell lines) or 48 h (patient ascites) to determine the induction of RS and intrinsic HRR status of cells/cultures. The impact of VE-821 (1 μM), PF-477736 (50 nM) and MK-1775 (100 nM) on RS and rucaparib-induced HRR was measured after 24/ 48 h exposure to drug alone or in combination with rucaparib. Concentrations were selected from single agent cytotoxicity data, where these concentrations impacted cell survival, without excessive cytotoxicity. Coverslips were washed twice with PBS before fixation with ice cold methanol for a minimum of 1 hr at -20°C then washed with 0.2% Triton-X-100 in PBS (cell lines) or 0.2% Triton-X-100 in 120 mM potassium chloride, 20 mM sodium chloride, 10 mM Tris and 1 mM EDTA, pH 8.0 (patient ascites). They were incubated in blocking buffer (2% BSA, 10% skimmed milk powder (Marvel, UK) and 10 % goat serum dissolved in relevant washing buffer), for 1 hr at room temperature. Fresh blocking buffer containing 1:500 of anti-RAD51 antibody mAb (Abcam, Cambridge, UK, ab133534) was added for 16-18 hr at 4°C before washes and the addition of anti-phospho H2A.X mAb (Merck, Kenilworth, NJ, United States, clone JBW301) diluted 1:1000 in buffer (2% BSA in relevant wash buffer) for 1 hr at room temperature. Following further washes both secondary antibodies Alexa Fluor 488 (anti-rabbit) and Alexa Fluor 546 (anti-mouse) (Invitrogen, Waltham, MA, USA) diluted 1:1000 were added to coverslips for 1 hr at room temperature in dark conditions before coverslips were washed and exposed to 0.5 μg/ml DAPI solution prior to mounting coverslips onto microscope slides using Prolong Glass Antifade mountant (Thermofisher, Waltham, MA, USA). Cells were imaged using Leica DM6 wide-field microscope (Leica Microsystems, Wetzlar, Germany) and analysed using Image J software. RAD51 foci were counted in γH2AX-positive cells (cells with > mean γH2AX foci value of untreated controls) to exclude cells that did not enter S-phase with SSB/trapped PARP1 and therefore do not have replication lesions that induce HRR. In instances where background γH2AX levels were high (HRD UWB cells and HRD ascites) foci in all cells were counted.

### Cell cycle analysis

Exponentially growing IGROV-1, UWB+ B1 and UWB cells were seeded and left to adhere for 24 h before exposure to drug for 30 h. Cell cycle analysis was performed by flow cytometry and in the final hour of drug treatment 10 μM EdU was added to the media containing drug to accurately identify cells in S-phase using a Click-iT™ EdU flow cytometry assay kit (Thermofisher, Waltham, MA, USA). Cells were harvested and fixed by 4% paraformaldehyde (15 minutes, room temperature and dark conditions) before centrifugation (1,600 rpm, 5 minutes) and washes (1 % BSA/PBS). Following further centrifugation (1,600 rpm, 5 minutes), Click-iT ™ reaction mixture was prepared according to the manufacturer’s instructions. Samples were incubated with reaction mixture then centrifuged (1,600 rpm, 5 minutes) and washed further (1% BSA/PBS). The cell pellet was resuspended in 1% BSA/PBS and 0.5 mg/ ml DAPI, diluted 1:250 (Thermofisher, Waltham, MA, USA) and analysis was performed using Attune Nxt Cytometer (Thermofisher, Waltham, MA, USA). De Novo FCS Express 7 software was used to analyse the data.

### Culturing patient ascites

Malignant ascites were collected from patients with ovarian cancer undergoing surgery or palliative drainage of ascites at the Queen Elizabeth Hospital, Gateshead, UK. Ethical approval and written consent was attained from North East Newcastle and North Tyneside 1 Research Ethics Committee for the collection of both clinical material and patient data. Samples were handled in accordance with the Human Tissue Act (2004). Ascites samples were initially processed by mixing ascites fluid 1:1 with RPMI-1640 medium, containing 20 % FBS, 1% penicillin/ streptomycin and transferred to culture flasks. Cultures were incubated at 37° C, 5% CO_2_ and 95% humidity, with medium replaced every 3-5 days, until cells had adhered and reached 70-80 % confluence.

### Growth inhibition by SRB assay in patient samples

Malignant ascites were established in culture and at first passage seeded at 500-1000 cells/well in 96-well plates, allowing 2 plates for drug treatments and 1 row of cells in a separate plate for the day 0 and allowed to adhere for 24 h. Medium was then replaced with media containing vehicle alone (DMSO control), rucaparib (0-10 μM) alone or in combination with VE-821 (1 μM), PF-477736 (50 nM) or MK-1775 (100 nM) in a final concentration of 0.5% DMSO and the day 0 row of cells was fixed with 25 μl/well methanol: acetic acid 3:1, washed dried and stored at 4°C. Once control cells reached 70-80% confluence they were fixed alongside treated cells, which were analysed alongside day 0 plates. Following washes with diH_2_O, 0.4% SRB solution in 1% acetic acid was added for 30 minutes (100 μl/well). After further washing 10 mM Tris buffer was added (pH 10.5, 100 μl/well) for 20 minutes on a plate shaker to solubilise the stain, prior to plates being read on the Omega FLUOstar (570 nm absorbance). The day 0 values were used to assess cell doubling times and growth inhibition was calculated by comparison between treated cells and vehicle control.

## Results

### Rucaparib-induced cytotoxicity, RS and activation of ATR, CHK1 and WEE1

The relative sensitivities of the cell lines to rucaparib cytotoxicity were measured by colony formation assays (Fig. 1A). As previously reported, rucaparib was more cytotoxic to HRD cells compared to matched HRP cells. The *BRCA2*-mutant V-C8 cells were >1,000-fold more sensitive than *BRCA2*-corrected V-C8.B2 cells (LC_50_ <0.01 μM vs >10 μM, p<0.001) and *BRCA1* mutant UWB were 62-fold more sensitive than *BRCA1*-corrected UWB +B1 cells (LC_50_: 0.19 μM vs 11.5 μM, p= 0.003). The other HRP cells in the panel displayed a narrow spectrum of sensitivity to rucaparib (LC_50_s: 8.0 μM to 12.3 μM, Fig. 1A; Supplementary Fig. S1A).

**Figure 1.**
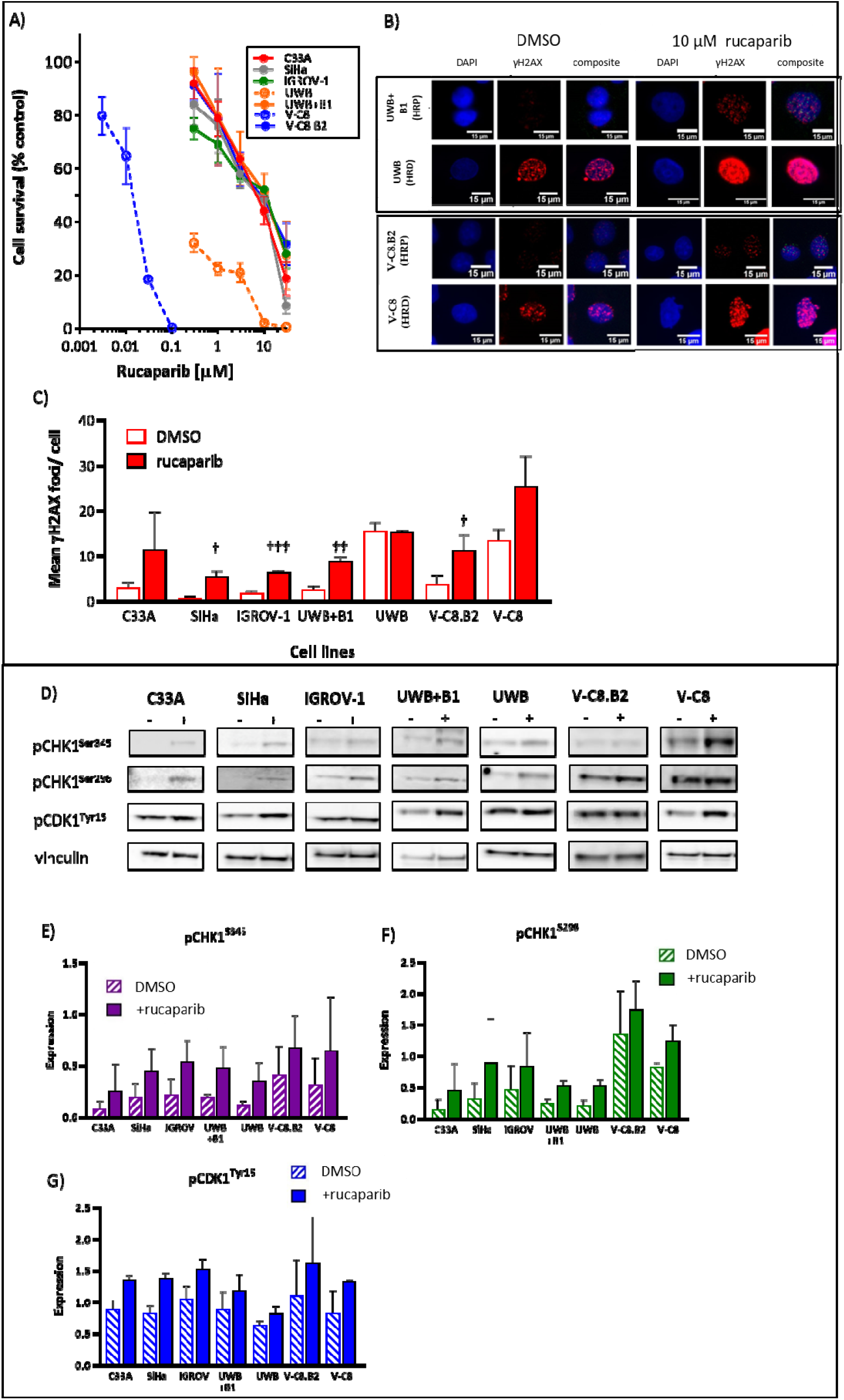
Rucaparib exerts differential cytotoxicity, induces replication stress (RS) and activates ATR, CHK1 and WEE1 in the cell panel. **A**. Colony formation-based cell survival following exposure of cells to the indicated concentration of rucaparib for 24 h. Data, percent survival relative to vehicle (DMSO) control, are mean ± SEM of 3 independent experiments. **B**. Induction of RS following exposure of cells to 10 μM rucaparib for 24 h, measured by γH2AX foci/cell. Representative images of paired cell lines are shown. **C**. Mean data from 3 independent experiments of the type shown in Figure 1B, significant induction by rucaparib is shown by † p<0.05, † † p<0.01, † † † p<0.0001 (scatter plots in supplementary 1C). **D**. Induction of phosphorylation of CHK1 S345, CHK1 S296 and CDK1 Y15 following exposure to 10 μM rucaparib for 24 h determined by Western blotting. Representative blots are shown. **E-G**. Pooled data for all cell lines from 3 independent experiments of the type shown in 1D. Each bar represents mean and standard error of expression normalised to vinculin loading control.

To determine if rucaparib-induced RS varied between cells we exposed them to 10 μM rucaparib for 24 h and measured RS by γH2AX foci (Fig. 1B,C). γH2AX foci typically detect DSB and collapsed replication forks, however under these conditions, the vast majority reflect the RS-induced DSBs caused by unrepaired SSBs and PARP1 trapped on the DNA meeting the replication machinery. γH2AX foci correlate well with pRPA foci, which is key in response to RS and is responsible for the regulation of replication fork restart and new origin firing (Supplementary Fig. S1B, 29). Rucaparib caused a 3 to 10-fold increase in γH2AX focus formation in HRP cells (Fig. 1C; Supplementary Fig. S1C), Basal levels of RS were significantly higher in *BRCA* mutant UWB (p=0.01) and V-C8 cells (p=0.04), compared to their matched HRP cells. Although rucaparib caused >2-fold increase in HRD V-C8 cells no increase was observed in HRD UWB cells.

To establish if the rucaparib-induced RS activated ATR, CHK1 and WEE1, we measured their phosphorylated targets (pCHK1S345, pCHK1S296 and pCDK1Y15, respectively) following exposure to rucaparib for 24 h. It is important to note that whereas pCHK1S345 and pCHK1S296 are relatively specific indicators of ATR and CHK1, respectively, increased pCDK1Y15 results from a combination of direct phosphorylation by WEE1 combined with inhibition of CDC25C dephosphorylation when CHK1 is activated (29). Rucaparib caused a 1.6 to 3.3-fold increase in pCHK1S345 and 1.2 to 3.1-fold increase in pCHK1S296 (Fig. 1D-F). Basal levels of CDK1 phosphorylation were high, possibly reflecting endogenous activity of MYT1 and WEE1, and, as a result, the induction by rucaparib was modest (1.3 to 1.7-fold) (Fig. 1 G). Surprisingly, despite high basal levels of RS in HRD UWB cells with no/modest further induction by rucaparib, the UWB cells did not have particularly high background levels of these phosphorylation targets and phosphorylation was increased by rucaparib to a similar extent as in HRP cells.

### ATR-CHK1 pathway activation by rucaparib is inhibited by VE-821, PF-477736 and MK-1775

To determine the effect of checkpoint kinase inhibitors on phosphorylated targets in cells exposed to PARPi, we treated cells with combinations of rucaparib (10 μM) and 1 μM VE-821 (ATRi), 50 nM/200 nM PF-477736 (CHK1i) or 100 nM/300 nM MK-1775 (WEE1i) for 24 h (Fig. 2A). VE-821 inhibited rucaparib-induced CHK1S345 phosphorylation by 64.4% to >100.0% (with inhibition >100% signifying inhibition of endogenous phosphorylation as well as that induced by rucaparib). VE-821 also inhibited downstream CHK1S296 phosphorylation by 38.1% to >100% and CDK1Y15 phosphorylation by 49.3% to >100.0% (Fig. 2B). PF-477736 inhibited rucaparib-induced CHK1S296 phosphorylation 76.5% to >100% in all cell lines, and inhibited CDK1Y15 phosphorylation 48.5% to >100% (Fig. 2C). Exposure to PF-477736 also caused a 5.6 to 31.3-fold increase in phosphorylated CHK1S345 upstream, indicating feedback activation of ATR when CHK1 is inhibited. MK-1775 reduced CDK1Y15 phosphorylation 79.8% to >100% (Fig. 2D). Additionally, upstream CHK1S296 phosphorylation was increased 1.2 to 8.5-fold and CHK1S345 phosphorylation increased 3.1 to 19.6-fold, further suggesting feedback mechanisms within the signalling cascade. Previously 1 μM VE-821, 50 nM PF-477736 and 100 nM MK-1775 inhibited cisplatin-induced activation of pCHK1S345, pCHK1S296 and pCDK1Y15 in cervical C33A an SiHa cells 79 to 93 %, 85 to 100 % and 54 to 58 %, respectively (30).

**Figure 2.**
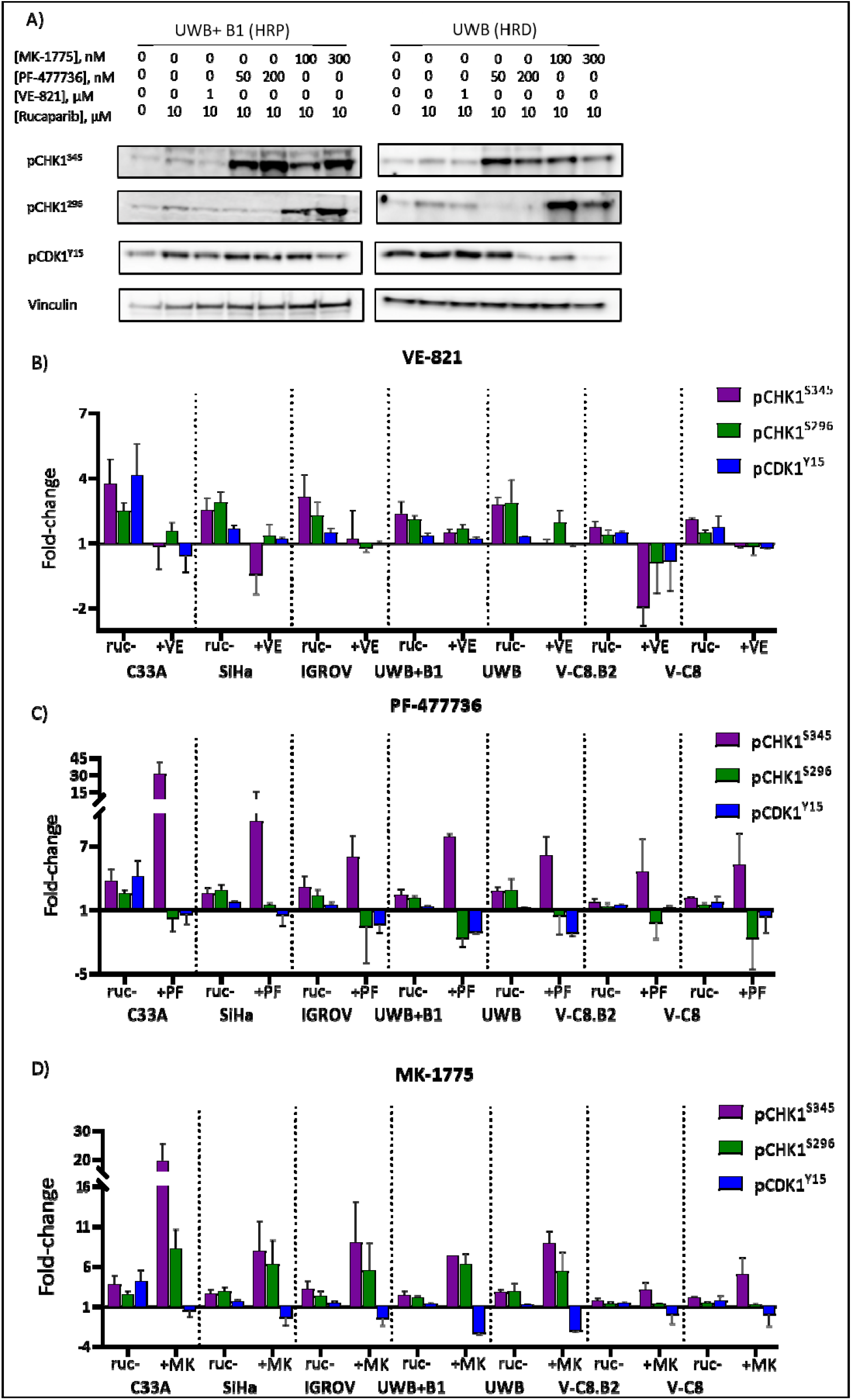
ATR-CHK1 pathway activation by rucaparib is inhibited by VE-821, PF-477736 and MK-1775. **A**. Cells were exposed to DMSO, 10 μM rucaparib alone or in the presence of 1 μM VE-821, 50 nM PF-477736, 200 nM PF-477736, 100 nM MK-1775 or 300 nM MK-1775 for 24 h prior to lysis. Representative Western blot of UWB1.289 and UWB1.298+B1 are shown **B-D**. Levels of pCHK1 S345, pCHK1 S296 and pCDK1 Y15 were measured by densitometry and normalised to relative vinculin loading control. Fold-change in expression (determined by densitometry) from DMSO control cells to cells treated with rucaparib +/- kinase inhibitors. Data shown are mean and standard error of 3 independent experiments.

### Single agent VE-821, PF-477736 and MK-1775 induced cytotoxicity

Cells displayed a spectrum of sensitivity to VE-821, PF-477736 and MK-1775 cytotoxicity. LC_50_ values varied 3.5-fold with VE-821 (0.6 to 2.1 μM), 5-fold with PF-477736 (22.4 to 125.5 nM) and 5-fold with MK-1775 (121.4 to 454.5 nM) (Fig. 2 E-G, Supplementary Fig. S1D). *BRCA1-*mutant HRD UWB cells were significantly more sensitive to the inhibitors than matched *BRCA1-*corrected HRP cells: ATRi VE-821 (LC_50_: 2.7-fold lower, p=0.05), CHK1i PF-477736 (LC_50_: 1.7-fold lower, p=0.01) and WEE1i MK-1775 (cell survival at 400 nM: 3.5-fold lower, p=0.05). However, *BRCA2-*mutated HRD V-C8 cells were no more sensitive to any of the checkpoint kinase inhibitors than their matched BRCA2-corrected HRP cells. Cells ranked in the same order of sensitivity for VE-821 and MK-1775 (most to least sensitive: UWB, C33A, UWB+B1, IGROV-1, V-C8, V-C8.B2, SiHa), but differed with PF-477736 (most to least sensitive: UWB, IGROV-1, UWB+B1, SiHa, V-C8, V-C8.B2, C33A). HPV negative C33A cells were more sensitive to VE-821 and MK-1775 compared to HPV positive SiHa cells, although C33A cells were the most resistant cell line to PF-477736. Concentrations (1 μM VE-821, 50 nM PF-477736 and 100 nM MK-1775) that were not excessively cytotoxic (cell kill <50% in HRP cells) across the panel of cells were selected for further study (Supplementary Fig. S1E).

### VE-821, PF-477736 and MK-1775 sensitised HRP cells to rucaparib but not matched HRD cells

To assess the interaction between PARP inhibition and ATR, CHK1 and WEE1 inhibition we determined the potentiation of rucaparib cytotoxicity by VE-821 (1 μM), PF-477736 (50 nM) and MK-1775 (100 nM) by colony formation assays (Fig. 3). Interestingly, ATR, CHK1 and WEE1 inhibitors each caused a significant rucaparib cytotoxicity sensitisation in HRP UWB +B1 (Two-way ANOVA, p<0.0001, p<0.0001 and p=0.002, respectively) and V-C8.B2 cells (p<0.0001), but no sensitisation in their HRD matched cells (Fig. 3A). Indeed, in UWB+B1 cells at 3 μM rucaparib the addition of VE-821 caused 2.6-fold sensitisation, such that cell survival was not significantly different to HRD UWB cells exposed to 3 μM rucaparib as a single agent, indicating resistance to rucaparib is completely overcome in UWB+B1 cells. VE-821, PF-477736 and MK-1775 also significantly sensitised the other HRP ovarian IGROV-1 (p<0.0001) and cervical C33A (p<0.0001) and SiHa (p=0.02, p=0.0005 and p=0.0023, respectively) cancer cells (Fig. 3B-D). Analysis of AUC of the cytotoxicity curves confirmed the significantly sensitisation of HRP cells to rucaparib (Fig. 3E). Whilst the extent of sensitisation varied between cell lines, VE-821, PF-477736 and MK-1775 were similar in the degree of sensitisation in each cell line they conferred: C33A (2.5 to 3.9-fold), SiHa (1.9 to 3.5-fold), IGROV-1 (1.6 to 2.1-fold), UWB+ B1 (1.1 to 2.1-fold) and V-C8.B2 (3.0 to 5.1-fold) with no inhibitor proving to be superior to the others (Supplementary Fig. S2).

**Figure 3.**
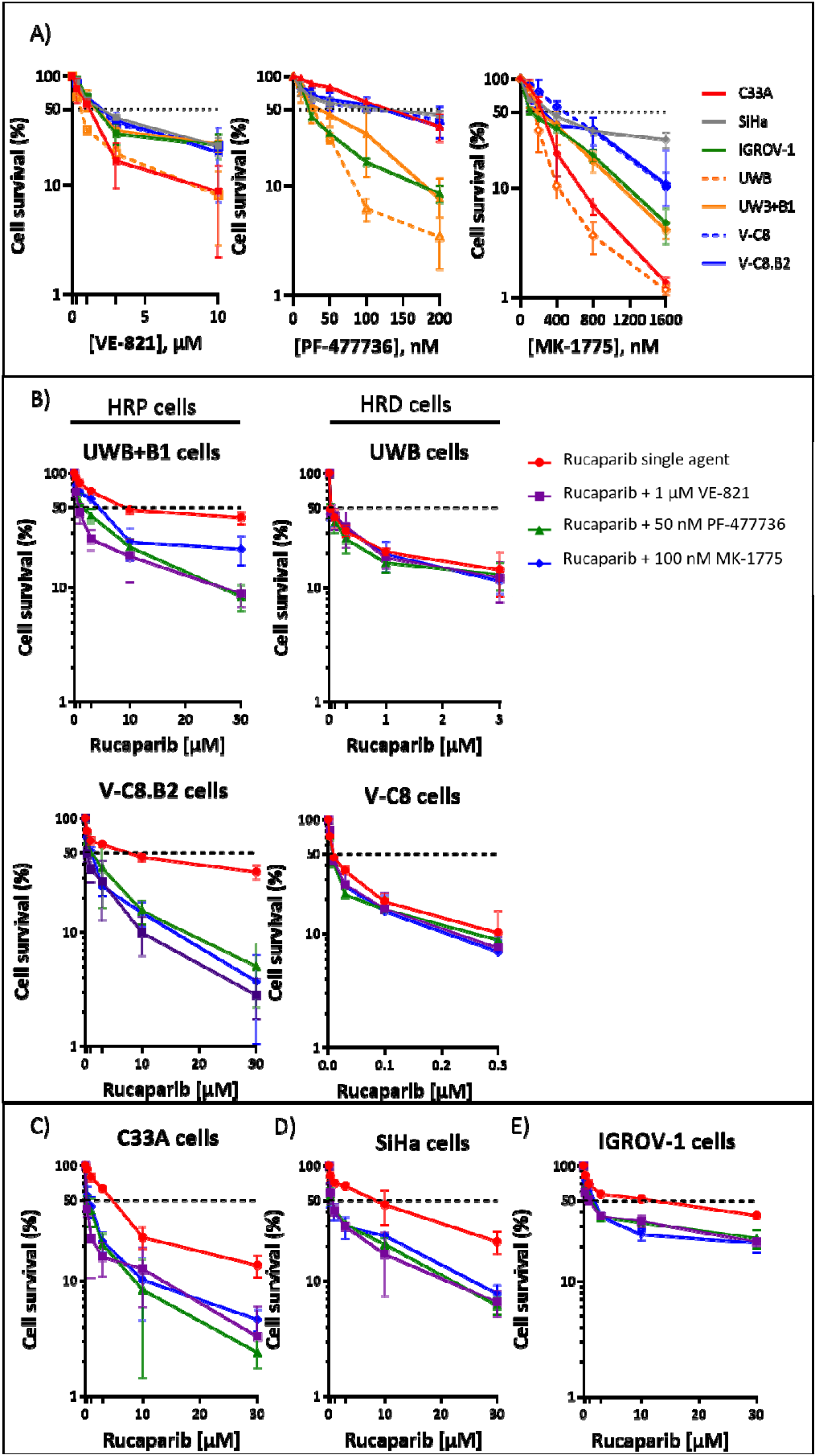
VE-821, PF-477736 and MK-1775 sensitised HRP cells to rucaparib but not matched HRD cells. **A**. Single agent cytotoxicity of VE-821, PF-477736 and MK-1775 (24 h exposure) was measured by colony formation assay. Data are mean ± SEM of 3 independent experiments. LC50 values and further cell survival data are displayed in supplementary 1D. **B-D**. Cells were exposed to rucaparib at indicated concentrations either as single agent or with the addition of 1 μM VE-821, 50 nM PF-477736 or 100 nM MK-1775 for 24 h prior to replacement with drug-free medium for 8-12 days to allow colony formation. Cell survival following exposure to combination treatments were normalised to single agent concentrations of VE-821 (1 μM), PF-477736 (50 nM) or MK-1775 (100 nM). Data are mean ± SEM of 3 independent experiments. **B**. matched HRP and HRD cells **C.** C33A cells **D.** SiHa cells and E. IGROV-1 cells.

### Rucaparib causes S-phase accumulation, which is abrogated by VE-821, PF-477736 and MK-1775

To determine the role of activation and inhibition of S and G2 cell cycle checkpoints in the potentiation of PARPi cytotoxicity by VE-821, PF-477736 and MK-1775, we performed cell cycle analysis in IGROV-1, UWB+ B1 and UWB cells (Fig. 4A, Supplementary Fig. S3). The induction of RS (as shown in Fig. 1B-D) by rucaparib (10 μM) caused an approximately 2-fold S-phase accumulation in all cells which was completely prevented by ATRi VE-821 (1 μM) co-exposure. CHK1i PF-477736 (50 nM) also blocked the rucaparib-induced S-phase accumulation in IGROV-1, UWB+ B1 and UWB cells (77.1 %, 97.8 % and 99.3 %, respectively) as did MK-1775 (100 nM) (64.0 %, 96.6 % and 80.0 %, respectively).

**Figure 4.**
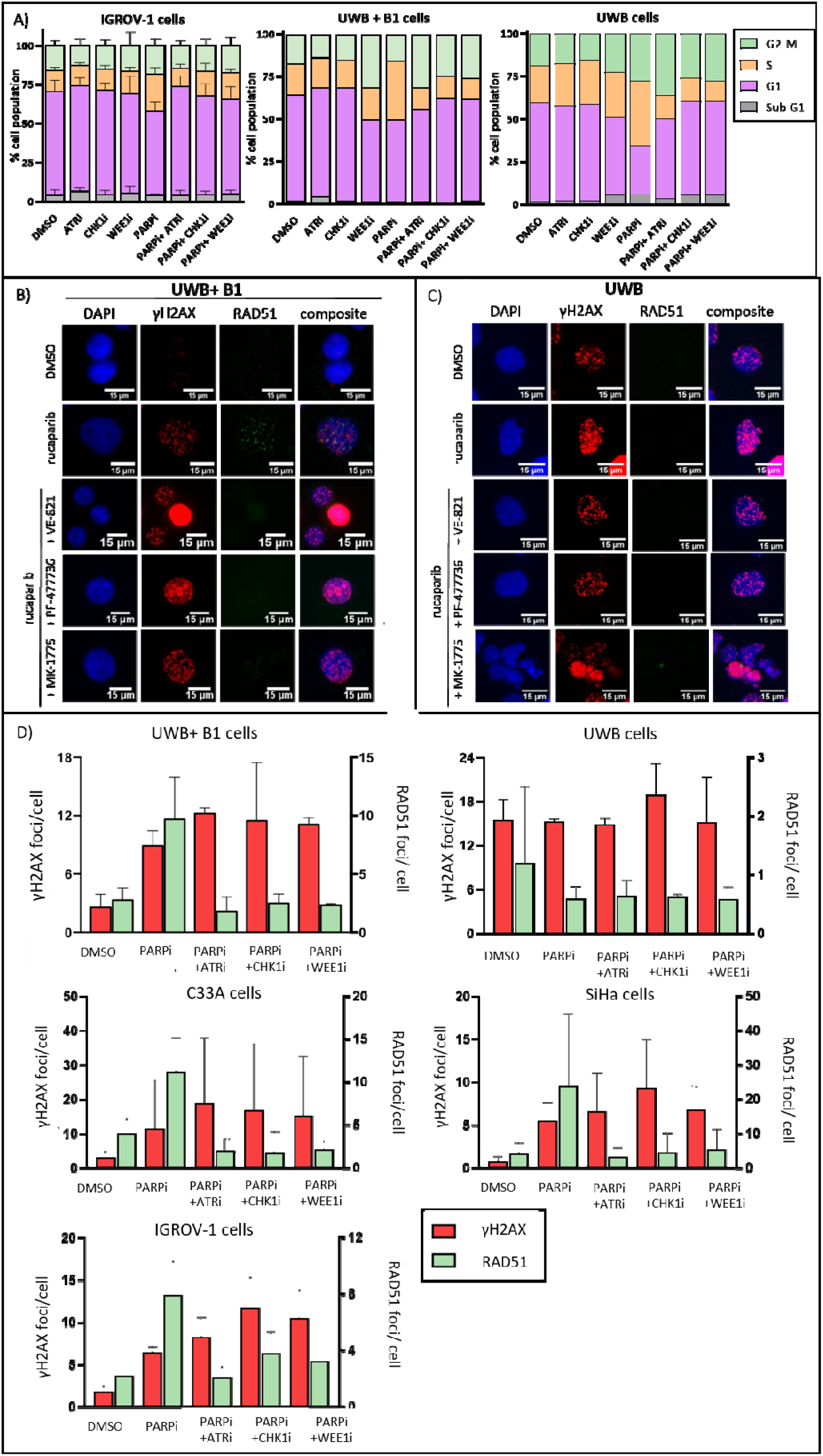
Impact of ATR, CHK1 and WEE1 inhibitors on cell cycle arrest and HRR induction by rucaparib. **A**. Cell cycle analysis of human ovarian cancer IGROV-1 and UWB paired cells following 24 h exposure to rucaparib single agent and in combination with VE-821 (1 μM), PF-477736 (50 nM) and MK-1775 (100 nM). 10 μM EdU was added to cells during the final hour of drug exposure. Data are mean and standard error of 3 independent experiments in IGROV-1 cells and mean of 2 independent experiments in UWB paired cells. Histograms and densitometry plots of representative experiments are shown in supplementary figure 3A-B. **B-C**. Formation of γH2AX foci was measured to indicate the induction of RS/DNA damage and RAD51 foci formation as an indicator of HRR function. Images are representative of observed differences between HRP and HRD cells upon exposure to DMSO, 10 μM rucaparib, 10 μM rucaparib+ 1 μM VE-821, 10 μM rucaparib + 50 nM PF-477736 or 10 μM rucaparib+ 100 nM MK-1775. **D**. Data show mean γH2AX foci/cell and mean RAD51 foci/γH2AX positive cell, with the exception of HRD UWB cells where RAD51 was analysed in all cells due to lack of induction of γH2AX with rucaparib. A total of 3 independent experiments are shown and standard error in cells treated with 10 μM rucaparib single agent and with the addition of 1 μM VE-821, 50 nM PF-477736 or 100 nM MK-1775 for 24 h prior to fixation.

### Rucaparib increases HRR activity, which is inhibited by VE-821, PF-477736 and MK-1775

The results from cell cycle analysis did not explain the observed differential sensitisation of HRP cells compared to HRD cells to rucaparib by the ATR, CHK1 and WEE1 inhibitors. Since ATR, CHK1 and WEE1 have also been implicated in HRR we investigated the impact of the inhibitors on HRR function as a possible explanation.

To assess HRR function we quantified RAD51 foci in cells exposed to rucaparib (10 μM) with or without VE-821 (1 μM), PF-477736 (50 nM) or MK-1775 (100 nM). Rucaparib caused a 5.4 ± 1.0-fold increase in RAD51 foci in γH2AX-positive HRP UWB+ B1 cells, compared to DMSO control cells (Fig. 4B,D), which was completely inhibited by VE-821 (>100%), and substantially reduced by PF-477736 (70%) and MK-1775 (87%). As predicted, there was no increase in RAD51 foci in *BRCA1* mutant UWB cells in response to rucaparib and this was not significantly changed by the checkpoint kinase inhibitors (Fig. 4C, D). Similar results were obtained in *BRCA2*-mutant and *BRCA2*-corrected V-C8 and V-C8.B2 cells (Supplementary Fig. S4B, Fig. S4C).

In the other cervical and ovarian HRP cell lines rucaparib increased RAD51 foci 4.8 to 5.4-fold, compared to DMSO control cells. VE-821, PF-477736 and MK-1775 each inhibited this HRR activity in C33A (>90%), SiHa (>95%) and IGROV-1 cells (65-90%) (Fig. 4D, Supplementary Fig. S4C). Since PARPi are synthetically lethal in HRD cells these data suggest that inhibition of HRR by the ATR, CHK1 and WEE1 inhibitors is likely to be the mechanism underpinning the differential potentiation of rucaparib cytotoxicity in HRP vs HRD matched cells, as the kinase inhibitors rendered HRP cells functionally HRD.

### Sensitisation and mechanistic observations translate to patient ascites cells

To determine if the sensitisation to rucaparib by VE-821, PF-477736 and MK-1775 observed in HRP cell lines was clinically relevant, we investigated the impact of the inhibitors on cell growth, RS and HRR in ascites cells from ovarian cancer patients.

In patient-derived ascites cells deemed HRP by RAD51 focus assay (PA225, PA233, PA240 and PA243), exposure to rucaparib (10 μM, 48 h) caused a 2.1 to 5.2-fold increase in γH2AX and a 2.4 to 4.6-fold increase in RAD51 foci (Fig. 5A). In HRD samples (PA236, PA241 and NC-16) basal levels of γH2AX foci were higher and only increased 1.2 to 2.2-fold by rucaparib (Fig. 5A), with no increase in RAD51 foci in these cells.

**Figure 5.**
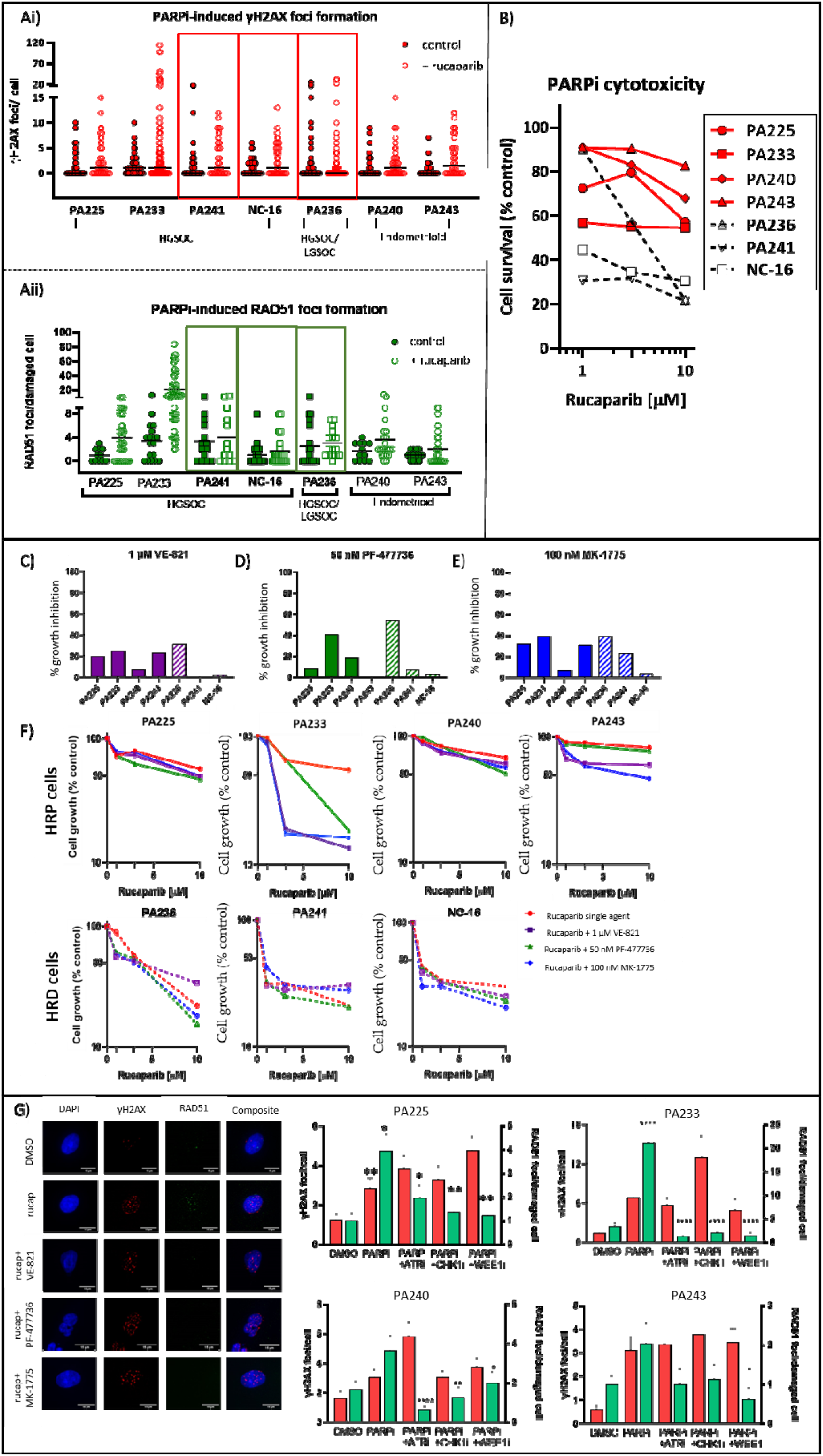
Impact of ATR, CHK1 and WEE1 inhibition on rucaparib induced cytotoxicity and HRR in patient ascites cells. **A**. γH2AX foci (i) and RAD51 foci (ii)were measured in ascites cells exposed to 10 μM rucaparib or vehicle (DMSO) control for 48 h, scatter plot with the mean bar shown. The highlighted samples are those identified as being HRD due to failure to induce RAD51 focus formation. **B**. Rucaparib induced growth inhibition measured by SRB staining intensity. Dashed lines represent HRD cells. Data are one experiment per primary culture. **C-E**. Growth inhibition by 1 μM VE-821, 50 nM PF-477736 and 100 nM MK-1775 determined by SRB staining intensity. **F**. Growth inhibition by rucaparib alone and in combination with VE-821 (1 μM), PF-477736 (50 nM) and MK-1775 (100 nM). Each graph represents one individual experiment per patient sample. **G**. Rucaparib-induction of HRR and inhibition by 1 μM VE-821, 50 nM PF-477736 and 100 nM MK-1775: the images shows PA240 as a representative HRP culture with the mean γH2AX foci/cell and RAD51/damaged cell.

SRB growth inhibition assays confirmed that HRD samples were more sensitive to rucaparib (GI_50_s: 0.6 to 3.6 μM) compared to HRP cultures, (GI_50_s: > 10 μM) (Fig. 5B; Supplementary Fig. S5). The concentrations of VE-821, PF-477736 and MK-1775 (1 μM, 50 nM and 100 nM, respectively) used in the previous experiments had only a modest effect on ascites cell growth as single agents (Fig. 5C-E). PA236 (HRD) was most susceptible to growth inhibition by VE-821, PF-477736 and MK-1775. However, the other HRD samples, PA241 and NC-16, were not more sensitive to any of the checkpoint kinase inhibitors, further suggesting that HRR status was not a determinant of sensitivity to these inhibitors.

Next, we determined the impact of VE-821 (1 μM), PF-477736 (50 nM) and MK-1775 (100 nM) on rucaparib-induced growth inhibition. In HRP ascites cells (PA225, PA233, PA40, PA243), VE-821, PF-477736 and MK-1775 sensitised cells to rucaparib but there was no sensitisation observed in the HRD cultures (Fig. 5F), replicating the observed differential sensitisation between HRP and HRD cell lines. Although there were differences in the extent of sensitisation between cultures, within each individual culture the inhibitors had a similar sensitising effect with no inhibitor proving superior to the others.

To investigate if the impact of VE-821, PF-477736 and MK-1775 on HRR function observed in cell lines was similar in the patient-derived cells we measured RAD51 foci in the ascites treated with rucaparib in the presence and absence of the inhibitors. As expected, the rucaparib-induction of RAD51 foci in HRP samples was reduced by VE-821, PF-477736 and MK-1775, and this reduction was significant in all HRP cultures except for PA243 (Fig. 5G; Supplementary Fig. S5).

## Discussion

Reversing the unresponsiveness of HRR proficient (HRP) cancers to PARPi therapy is an unmet clinical need. To address this, several clinical trials are investigating PARPi in combination with ATR, CHK1 and WEE1 inhibitors. In this study we investigated how inhibition of ATR, CHK1 and WEE1 sensitise HRP cells to PARPi therapy and which of the three checkpoint inhibitors would be most effective. We used paired HRP and HRD cells to determine the underlying mechanism was inhibition of cell cycle checkpoints or by impairing HRR function.

Both HRD cell lines (UWB and V-C8) were more sensitive to rucaparib compared to HRP matched counterparts as reported previously (14, 31). *BRCA2* mutation conferred greater sensitivity to rucaparib in V-C8 cells (>1000-fold), compared to a *BRCA1* mutation in UWB cells (62-fold). This may be because of BRCA1’s major role in end resection, which may not be as important for PARP-induced collapsed replication forks that would likely have single-stranded DNA (ssDNA) overhangs already. Under these circumstances BRCA2 may have a more critical role in mediating the recruitment of RAD51, to promote strand invasion and initiate synthesis in HRR (32, 33). Interestingly, results from the TRITON2 study (NCT02952534) showed clinical activity of rucaparib was greater in castrate-resistant prostate cancer patients with *BRCA2* mutations compared to *BRCA1* mutations (34). Further differences were observed in the sensitivity of cell lines to single agent ATR, CHK1 and WEE1 inhibitors, as HRD UWB cells were significantly more sensitive to all 3 kinase inhibitors compared to HRP UWB+B1 cells, however HRD V-C8 cells were no more sensitive to any of the inhibitors. This could suggest that clinically, although *BRCA1* mutation could be a predictive biomarker for ATR/CHK1/WEE1 inhibitor sensitivity, more broadly HRR status may not be useful and warrants further investigation. Caveats to these hypotheses are that the UWB cells were derived from a *BRCA1* mutant tumour that presumably adapted by engaging other DDR mechanisms to promote its vigorous survival, whereas in contrast, V-C8 cells were isolated following mutagenesis of their parental wildtype V79 cells (35) and are hence less likely to have evolved adaptive mechanisms. The different species origin of these cells may also affect their response.

RS results in γH2AX foci and signalling via ATR, CHK1 and WEE1. However, the higher levels of endogenous RS in HRD cells was not reflected in downstream phosphorylation events. Interestingly, HRP cervical C33A cells were the most sensitive to ATRi VE-821 and WEE1i MK-1775 and these cells also had the highest basal RS (γH2AX foci/cell) of the HRP cells. Similarly, HRD culture PA236 had the highest basal RS levels and this culture was also most sensitive to growth inhibition by VE-821, PF-477736 and MK-1775, further supporting RS as a potential biomarker for sensitivity to the kinase inhibitors (Fig. 5A, C). There was a general trend that suggested RS (measured by mean basal γH2AX foci/cell) was a determinant of sensitivity in cell lines to VE-821 and probably PF-477736 but not MK-1775 (Supplementary Fig. S6), further suggesting basal RS to be a biomarker for sensitivity to ATR, CHK1 and WEE1 inhibitors (36, 37, 38).

The data here suggest a potential role for HPV because HPV-negative C33A cells were most sensitive to both VE-821 and MK-1775, and HPV-positive SiHa cells were most resistant to VE-821 and MK-1775. However, this may not be universal as studies with a broader panel of cervical cancer cell lines suggest HPV does not impact on sensitivity to these agents, presumably because the non-HPV infected cells had pathological mutations in p53 and pRB (30).

The synergistic impact of PARPi with ATR, CHK1 or WEE1 inhibitors has been widely reported *in vitro* and *in vivo*. We previously demonstrated CHK1i PF-477736 sensitised rucaparib-treated HRP V-C8.B2 cells but not matched HRD V-C8 cells (14), suggesting HRR inhibition was the primary driving factor. Here, we broadened the use of cell cycle checkpoint kinase inhibitors to include ATRi VE-821 and WEE1i MK-1775 and discovered that all 3 inhibitors sensitised HRP cells, with no inhibitor proving superior to the others (Fig. 3). Our data showed that the rucaparib-induced RS activated the cell cycle checkpoint kinases (Fig. 1B-G), and that VE-821, PF-477736 and MK-1775 inhibited their relevant targets (Fig. 2A-D). This rucaparib-induced RS resulted in an increase in S-phase accumulation, which was reduced by all 3 kinase inhibitors to the same extent in HRP and HRD matched cells (Fig. 4). VE-821, PF-477736 and MK-1775 also equally inhibited rucaparib-induced HRR in HRP cells, frequently to below basal levels (Fig. 4D). Since the synergistic cytotoxicity of VE-821, PF-477736 and MK-1775 with rucaparib was only observed in HRP cells (Fig. 3), the most likely explanation is that the synergy is *via* inhibition of HRR causing the HRP cells to behave as HRD cells targetable by PARPi. In HRP cells there was a significant correlation with all 3 kinase inhibitors between the fold-sensitisation of cytotoxicity (Fig. 3E) and the inhibition of HRR (fold-reduction of RAD51 foci, Figure 4D), (Supplementary Fig. S7), defining a clear link between the degree of HRR impairment caused by the kinase inhibitors and the subsequent sensitisation of resistant HRP cells to rucaparib. Despite the comprehensive published evidence regarding the involvement of ATR, CHK1 and WEE1 in promoting HRR, it is difficult to reconcile the idea that ATR or CHK1 inhibition could be *directly* responsible for the induction of the HRD phenotype, as PF-477736 activated ATR activity and MK-17775 activated both CHK1 and ATR but all 3 inhibitors equally inhibited HRR. Data in this study showed ATRi VE-821 and CHK1i PF-477736 caused a similar reduction in pCDK1Y15 as WEE1i MK-1775 (Fig. 2B-D), suggesting that WEE1 or a downstream target of WEE1 may be ultimately responsible for promoting HRR.

Unquestionably, the key question to address is if these findings translate directly to the clinical situation. Our investigations in cultures of patient’s malignant ovarian cancer ascites suggest this is the case, as VE-821, PF-477736 and MK-1775 each blocked rucaparib-induced RAD51 focus formation in HRP patient cultures (Fig. 5G) and only sensitised HRP cultures to rucaparib (Figure 5F), strengthening the proposal that inhibition of HRR is the primary mechanism of action underpinning the synergy.

These findings support the notion that ATR, CHK1 and WEE1 inhibitors would sensitise tumours that are resistant to PARPi, by virtue of HRP genotype or restoration of HRR function in e.g. *BRCA* mutant tumours, clinically. However, such combinations may not be beneficial in those patients with HRD tumours and may only lead to an increase in toxicity. The real-time assessment of HRR function shown here by RAD51 focus formation in patient tumour material would be a good candidate for a predictive biomarker to identify HRD tumours, and further investigation of these methods has been recently recommended (39). Our data suggest its use could be expanded to establish whether patients would benefit most from combination of PARPi with ATR/CHK1/WEE1i (HRP tumours) or PARPi as a monotherapy (HRD tumours).

---

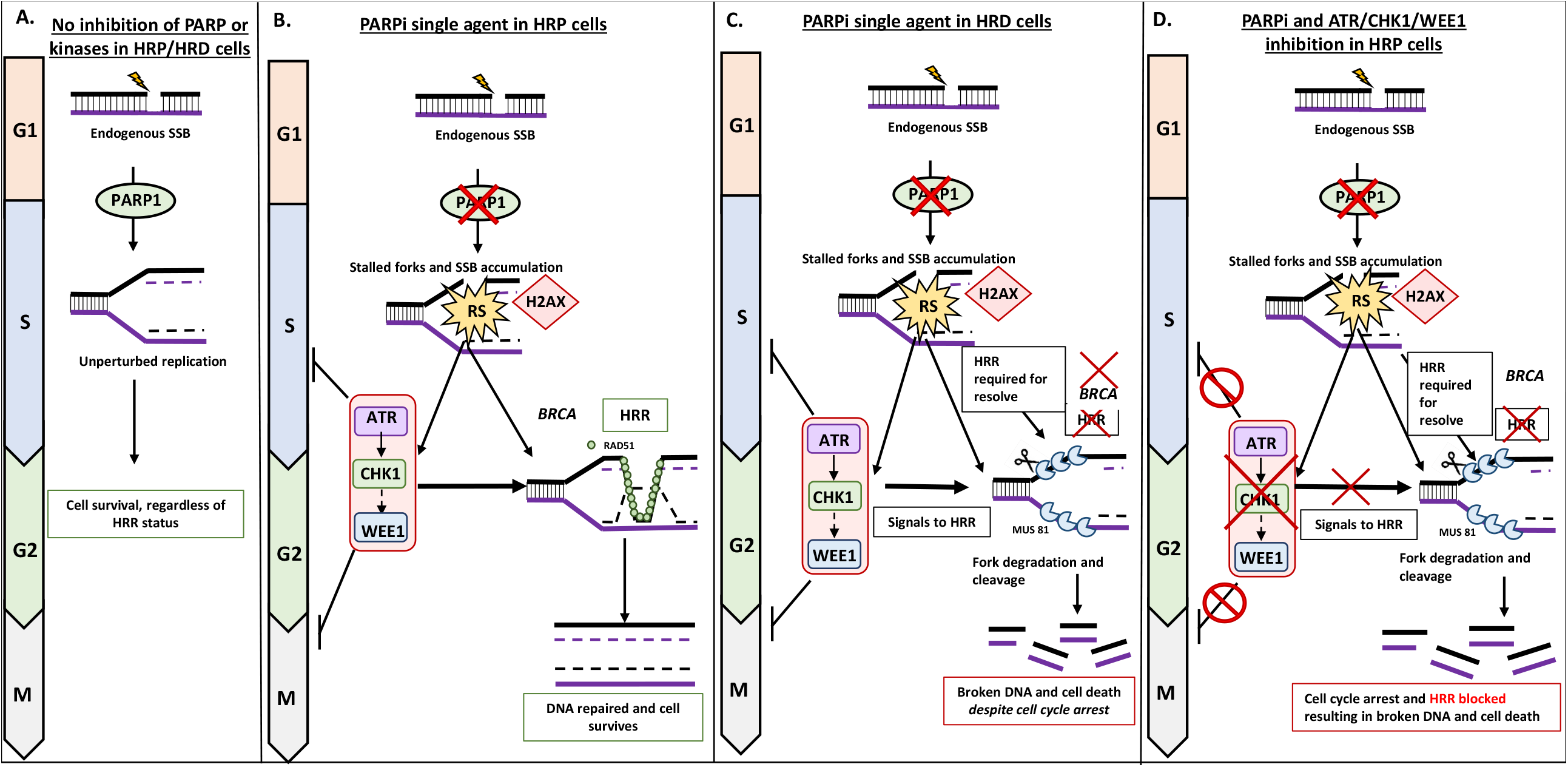
The proposed mechanism of action underpinning observed sensitisation of PARPi-treated cells by ATR, CHK1 and WEE1 inhibition. **A**. When an endogenous single-strand break (SSB) occurs in an HRP or HRD cell and PARP1 is present, the cell can repair via base excision repair, the first line of defence against DNA damage. The activation of PARP1 by the SSB subsequently leads to the recruitment of XRCC1, Pol β and Lig III to repair the DNA, resulting in unperturbed replication and cell survival. **B**. Exposure to PARPi results in the accumulation of unrepaired SSBs, resulting in stalled replication forks, causing replication stress (RS). The PARPi-induced RS signals to ATR, which initiates a signalling cascade to CHK1 and WEE1 downstream. This cascade is involved in halting the cell cycle, by preventing S-phase progression and entry into mitosis, allowing repair to occur. These replication-associated lesions rely on HRR for resolve, which ATR, CHK1 and WEE1 each signal too. In an HRP cell DNA is repaired and the cell survives, despite exposure to a PARPi. **C**. In HRD cells, which cannot rely on HRR for resolve of the PARPi-induced damage, fork degradation and cleavage occur, which ultimately results in cell death, despite PARPi-induced RS signalling to ATR, CHK1 and WEE1 for cell cycle arrest. **D**. When HRP cells are treated with single agent PARPi, the replication-associated lesions are resolved and the cell survives (seen in **B**.). However, when the HRP cell is also exposed to an ATR, CHK1 or WEE1 inhibitor HRR is inhibited in the HRP cell, rendering it sensitive to the PARPi. With ATR, CHK1 or WEE1 inhibited there is no cell cycle arrest and more importantly blocked HRR, resulting in broken DNA and ultimately cell death.

