## Supplementary material for "Overcoming PARP inhibitor resistance by inducing a homologous recombination repair defective phenotype with ATR, CHK1 and WEE1 inhibitors": Figures S1-S7

### Slide 1
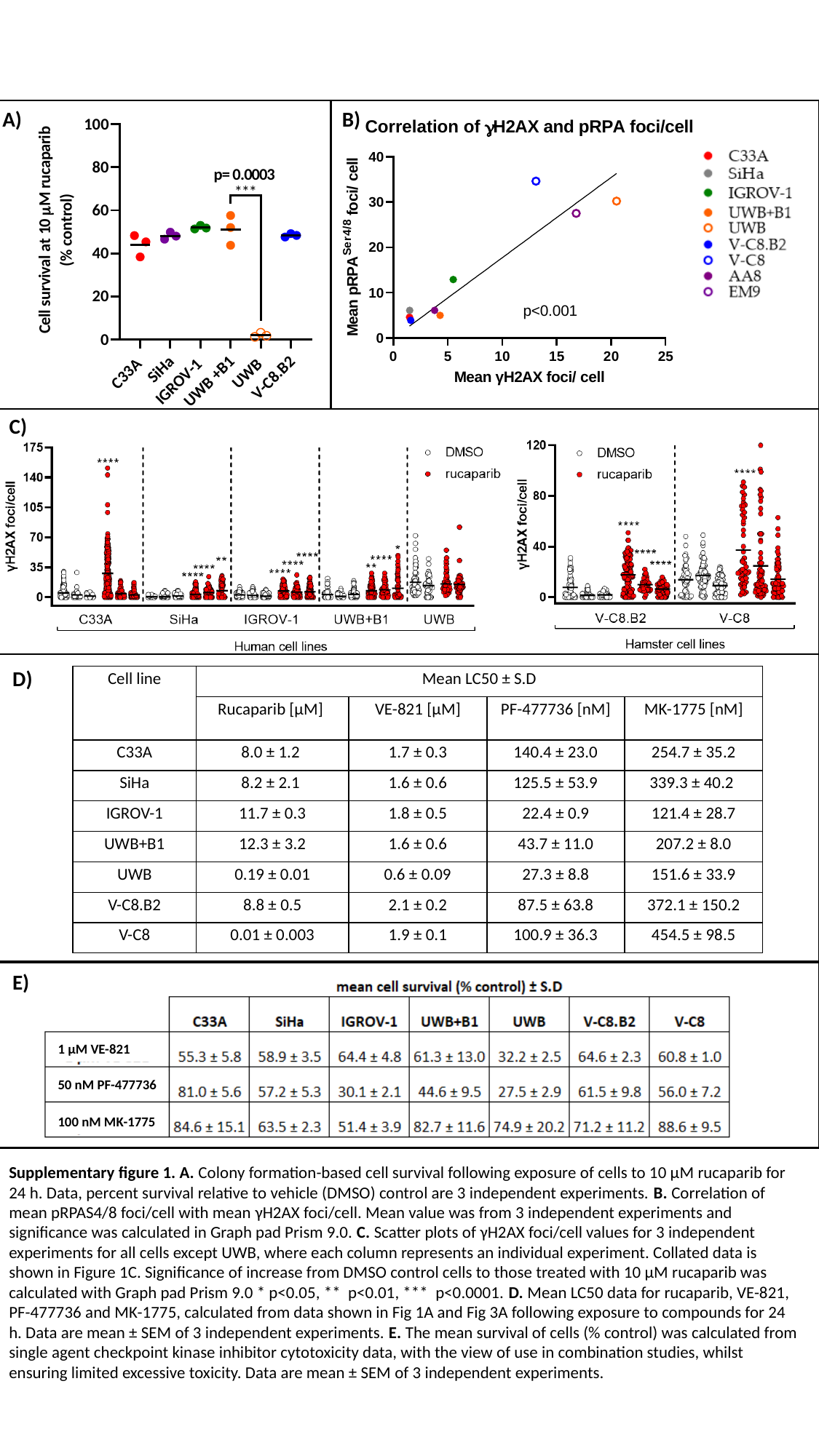

A)
B)
C)
D)
| Cell line | Mean LC50 ± S.D | | | |
| --- | --- | --- | --- | --- |
| | Rucaparib [µM] | VE-821 [µM] | PF-477736 [nM] | MK-1775 [nM] |
| C33A | 8.0 ± 1.2 | 1.7 ± 0.3 | 140.4 ± 23.0 | 254.7 ± 35.2 |
| SiHa | 8.2 ± 2.1 | 1.6 ± 0.6 | 125.5 ± 53.9 | 339.3 ± 40.2 |
| IGROV-1 | 11.7 ± 0.3 | 1.8 ± 0.5 | 22.4 ± 0.9 | 121.4 ± 28.7 |
| UWB+B1 | 12.3 ± 3.2 | 1.6 ± 0.6 | 43.7 ± 11.0 | 207.2 ± 8.0 |
| UWB | 0.19 ± 0.01 | 0.6 ± 0.09 | 27.3 ± 8.8 | 151.6 ± 33.9 |
| V-C8.B2 | 8.8 ± 0.5 | 2.1 ± 0.2 | 87.5 ± 63.8 | 372.1 ± 150.2 |
| V-C8 | 0.01 ± 0.003 | 1.9 ± 0.1 | 100.9 ± 36.3 | 454.5 ± 98.5 |
E)
1 µM VE-821
50 nM PF-477736
100 nM MK-1775
Supplementary figure 1. A. Colony formation-based cell survival following exposure of cells to 10 µM rucaparib for 24 h. Data, percent survival relative to vehicle (DMSO) control are 3 independent experiments. B. Correlation of mean pRPAS4/8 foci/cell with mean γH2AX foci/cell. Mean value was from 3 independent experiments and significance was calculated in Graph pad Prism 9.0. C. Scatter plots of γH2AX foci/cell values for 3 independent experiments for all cells except UWB, where each column represents an individual experiment. Collated data is shown in Figure 1C. Significance of increase from DMSO control cells to those treated with 10 µM rucaparib was calculated with Graph pad Prism 9.0 * p<0.05, ** p<0.01, *** p<0.0001. D. Mean LC50 data for rucaparib, VE-821, PF-477736 and MK-1775, calculated from data shown in Fig 1A and Fig 3A following exposure to compounds for 24 h. Data are mean ± SEM of 3 independent experiments. E. The mean survival of cells (% control) was calculated from single agent checkpoint kinase inhibitor cytotoxicity data, with the view of use in combination studies, whilst ensuring limited excessive toxicity. Data are mean ± SEM of 3 independent experiments.

### Slide 2
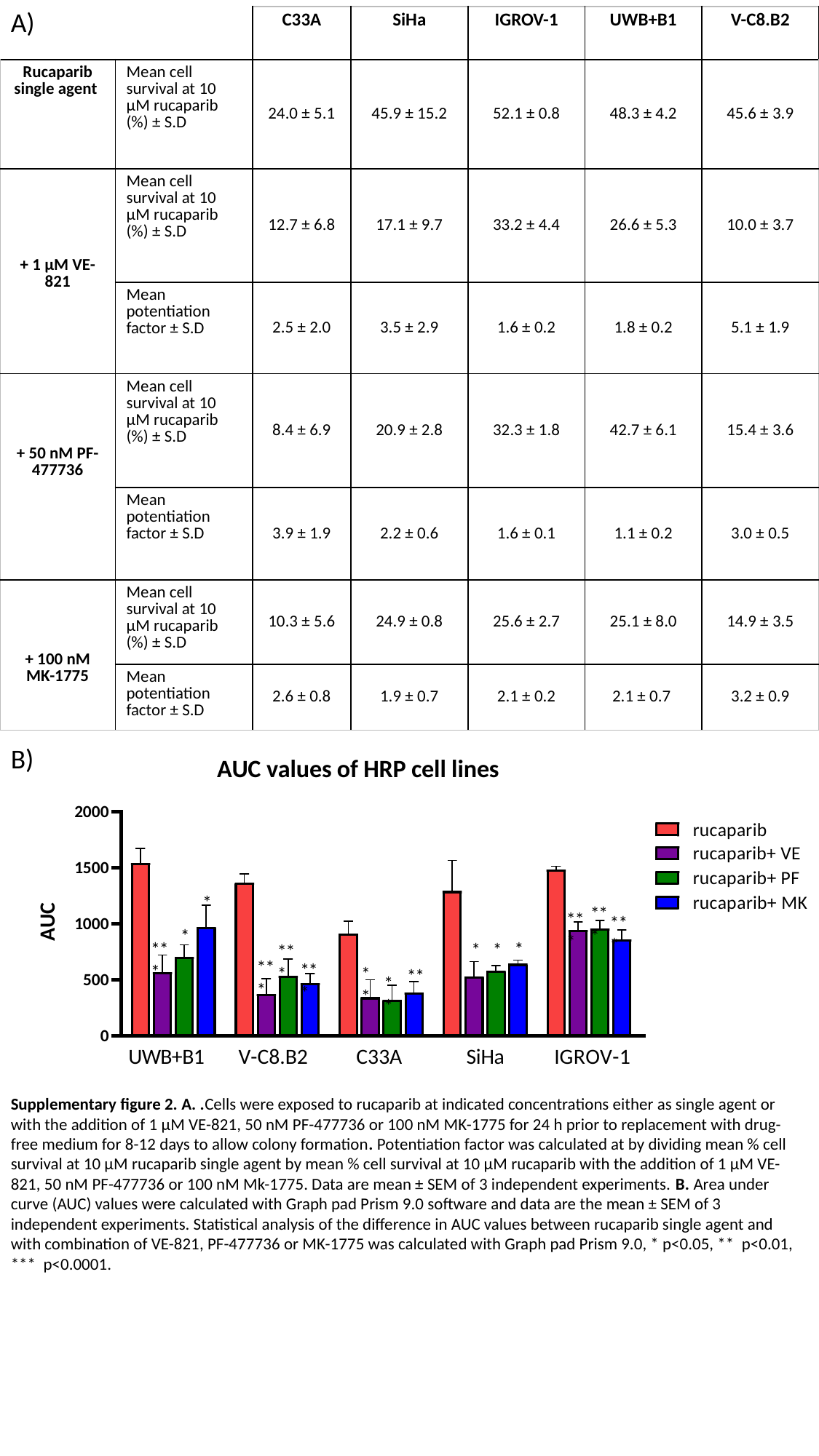

A)
| | | C33A | SiHa | IGROV-1 | UWB+B1 | V-C8.B2 |
| --- | --- | --- | --- | --- | --- | --- |
| Rucaparib single agent | Mean cell survival at 10 µM rucaparib (%) ± S.D | 24.0 ± 5.1 | 45.9 ± 15.2 | 52.1 ± 0.8 | 48.3 ± 4.2 | 45.6 ± 3.9 |
| + 1 µM VE-821 | Mean cell survival at 10 µM rucaparib (%) ± S.D | 12.7 ± 6.8 | 17.1 ± 9.7 | 33.2 ± 4.4 | 26.6 ± 5.3 | 10.0 ± 3.7 |
| | Mean potentiation factor ± S.D | 2.5 ± 2.0 | 3.5 ± 2.9 | 1.6 ± 0.2 | 1.8 ± 0.2 | 5.1 ± 1.9 |
| + 50 nM PF-477736 | Mean cell survival at 10 µM rucaparib (%) ± S.D | 8.4 ± 6.9 | 20.9 ± 2.8 | 32.3 ± 1.8 | 42.7 ± 6.1 | 15.4 ± 3.6 |
| | Mean potentiation factor ± S.D | 3.9 ± 1.9 | 2.2 ± 0.6 | 1.6 ± 0.1 | 1.1 ± 0.2 | 3.0 ± 0.5 |
| + 100 nM MK-1775 | Mean cell survival at 10 µM rucaparib (%) ± S.D | 10.3 ± 5.6 | 24.9 ± 0.8 | 25.6 ± 2.7 | 25.1 ± 8.0 | 14.9 ± 3.5 |
| | Mean potentiation factor ± S.D | 2.6 ± 0.8 | 1.9 ± 0.7 | 2.1 ± 0.2 | 2.1 ± 0.7 | 3.2 ± 0.9 |
B)
Supplementary figure 2. A. .Cells were exposed to rucaparib at indicated concentrations either as single agent or with the addition of 1 µM VE-821, 50 nM PF-477736 or 100 nM MK-1775 for 24 h prior to replacement with drug-free medium for 8-12 days to allow colony formation. Potentiation factor was calculated at by dividing mean % cell survival at 10 µM rucaparib single agent by mean % cell survival at 10 µM rucaparib with the addition of 1 µM VE-821, 50 nM PF-477736 or 100 nM Mk-1775. Data are mean ± SEM of 3 independent experiments. B. Area under curve (AUC) values were calculated with Graph pad Prism 9.0 software and data are the mean ± SEM of 3 independent experiments. Statistical analysis of the difference in AUC values between rucaparib single agent and with combination of VE-821, PF-477736 or MK-1775 was calculated with Graph pad Prism 9.0, * p<0.05, ** p<0.01, *** p<0.0001.

### Slide 3
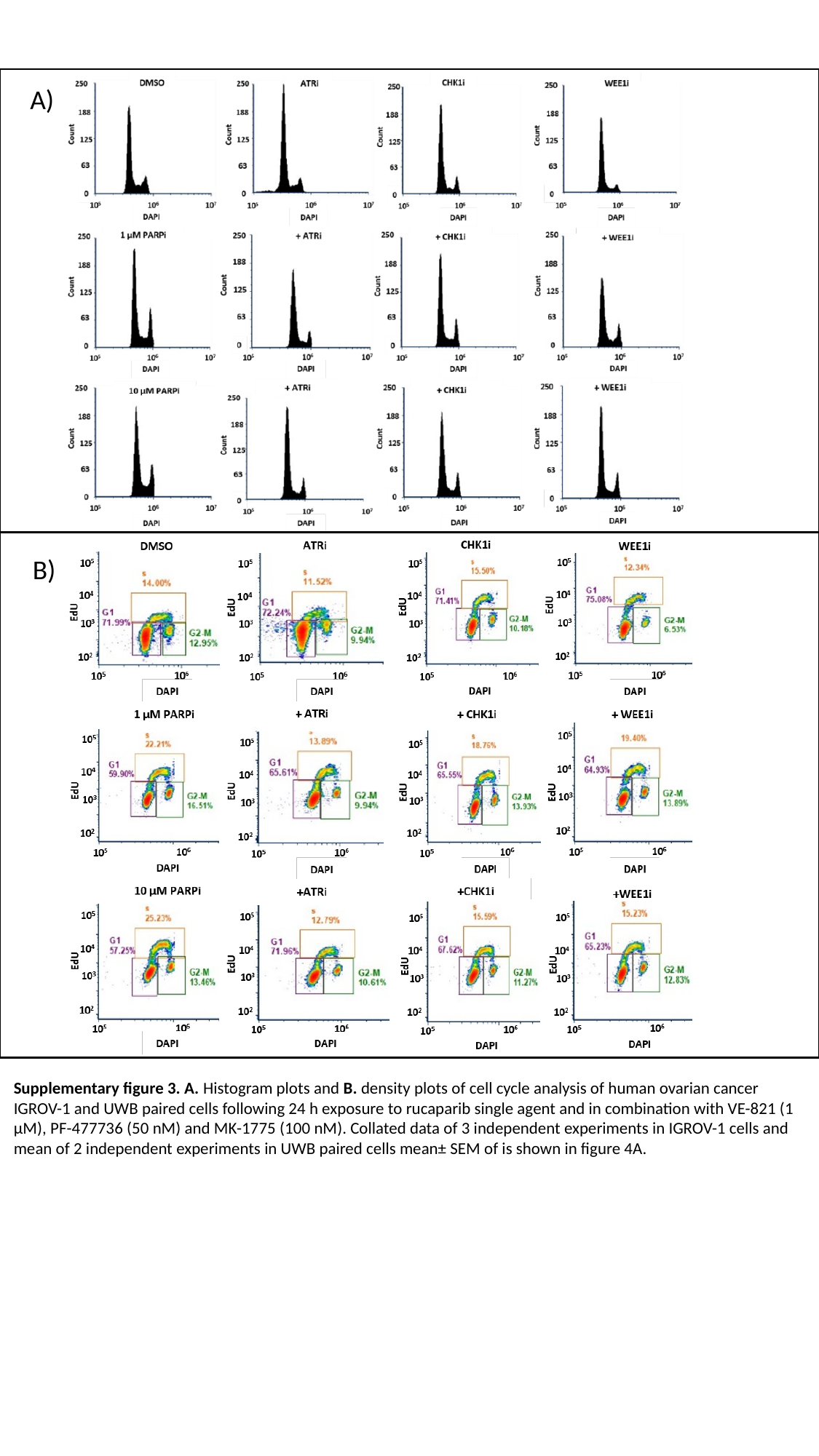

A)
B)
Supplementary figure 3. A. Histogram plots and B. density plots of cell cycle analysis of human ovarian cancer IGROV-1 and UWB paired cells following 24 h exposure to rucaparib single agent and in combination with VE-821 (1 µM), PF-477736 (50 nM) and MK-1775 (100 nM). Collated data of 3 independent experiments in IGROV-1 cells and mean of 2 independent experiments in UWB paired cells mean± SEM of is shown in figure 4A.

### Slide 4
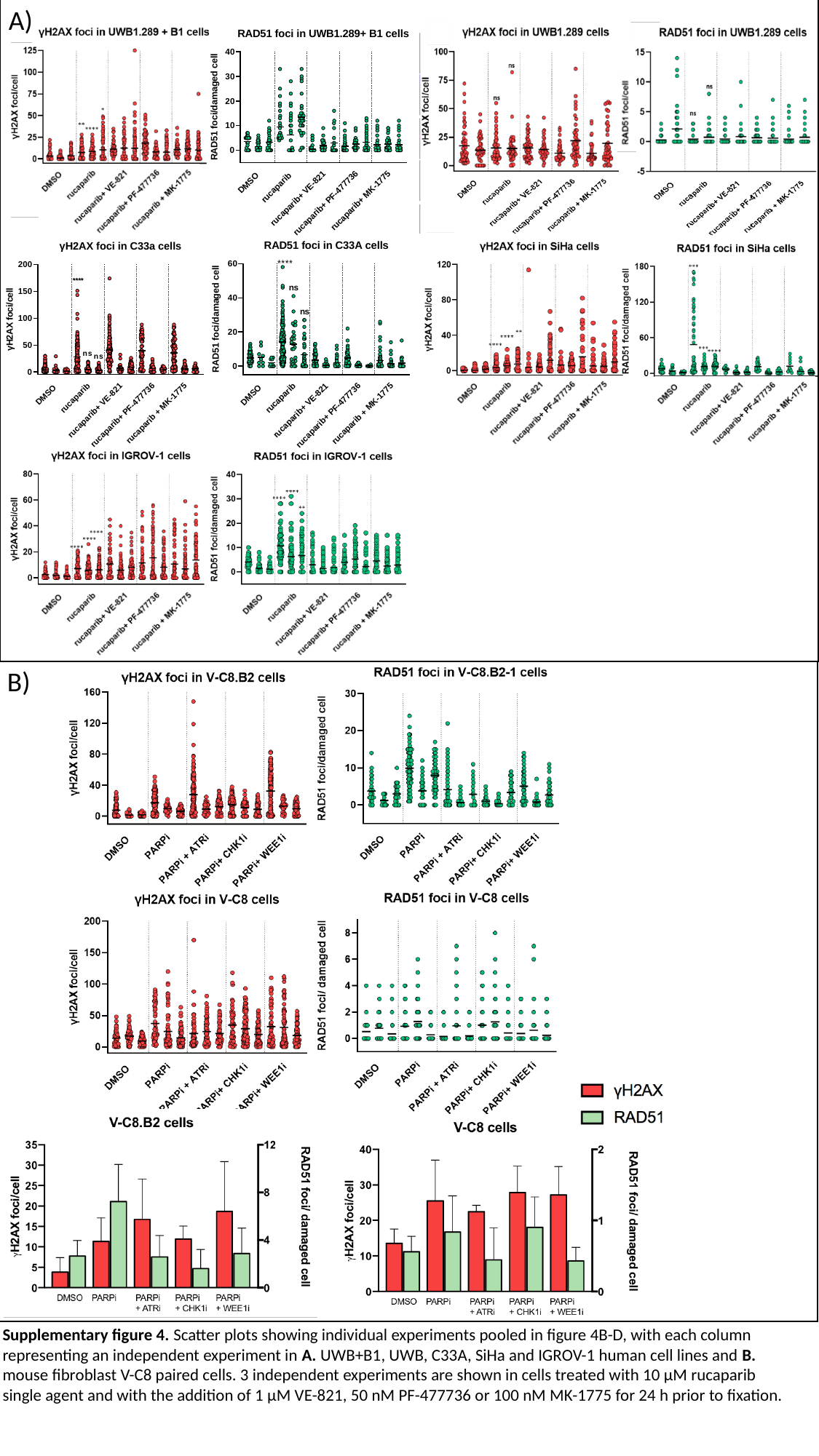

A)
B)
Supplementary figure 4. Scatter plots showing individual experiments pooled in figure 4B-D, with each column representing an independent experiment in A. UWB+B1, UWB, C33A, SiHa and IGROV-1 human cell lines and B. mouse fibroblast V-C8 paired cells. 3 independent experiments are shown in cells treated with 10 µM rucaparib single agent and with the addition of 1 µM VE-821, 50 nM PF-477736 or 100 nM MK-1775 for 24 h prior to fixation.

### Slide 5
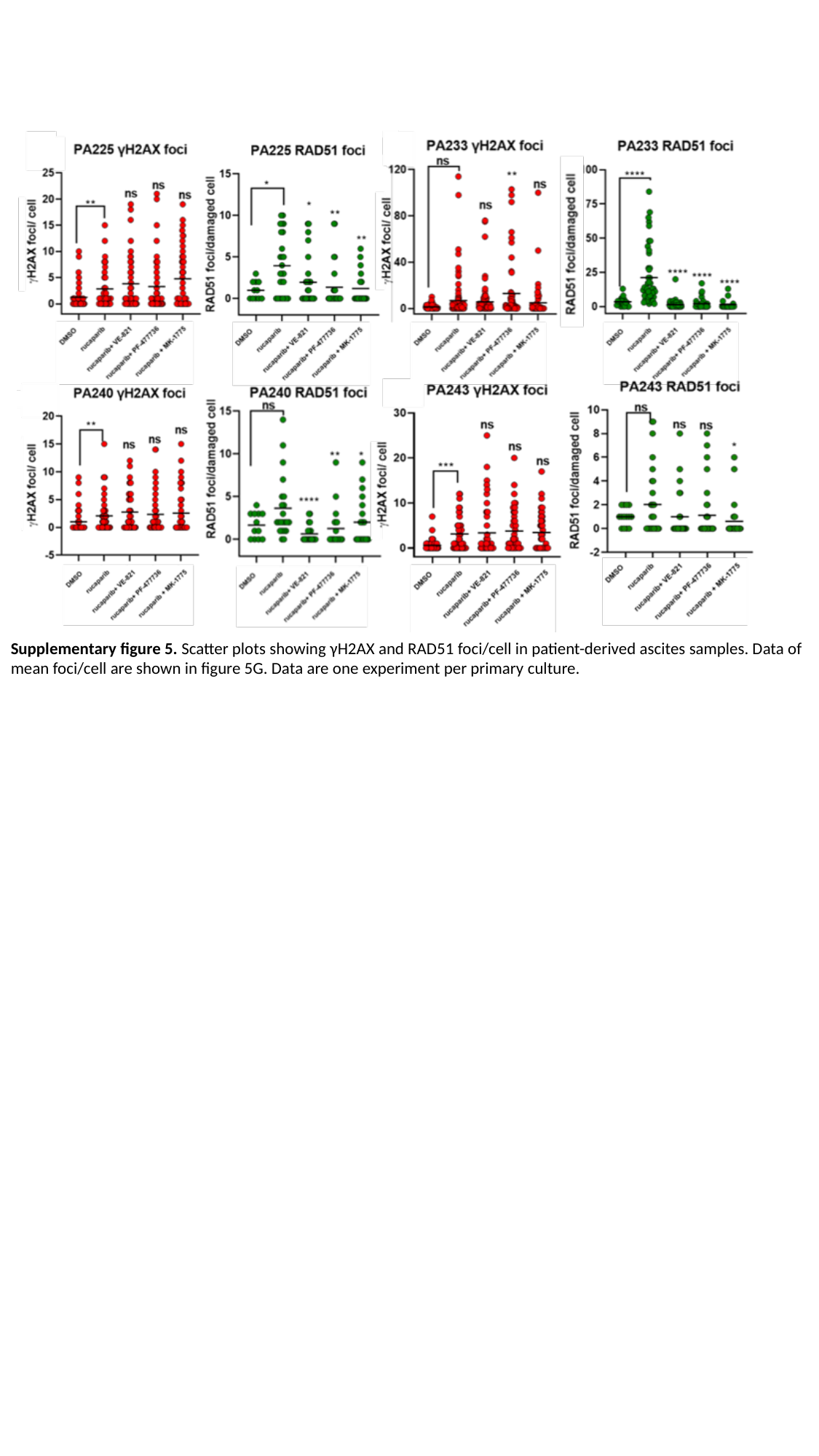

Supplementary figure 5. Scatter plots showing γH2AX and RAD51 foci/cell in patient-derived ascites samples. Data of mean foci/cell are shown in figure 5G. Data are one experiment per primary culture.

### Slide 6
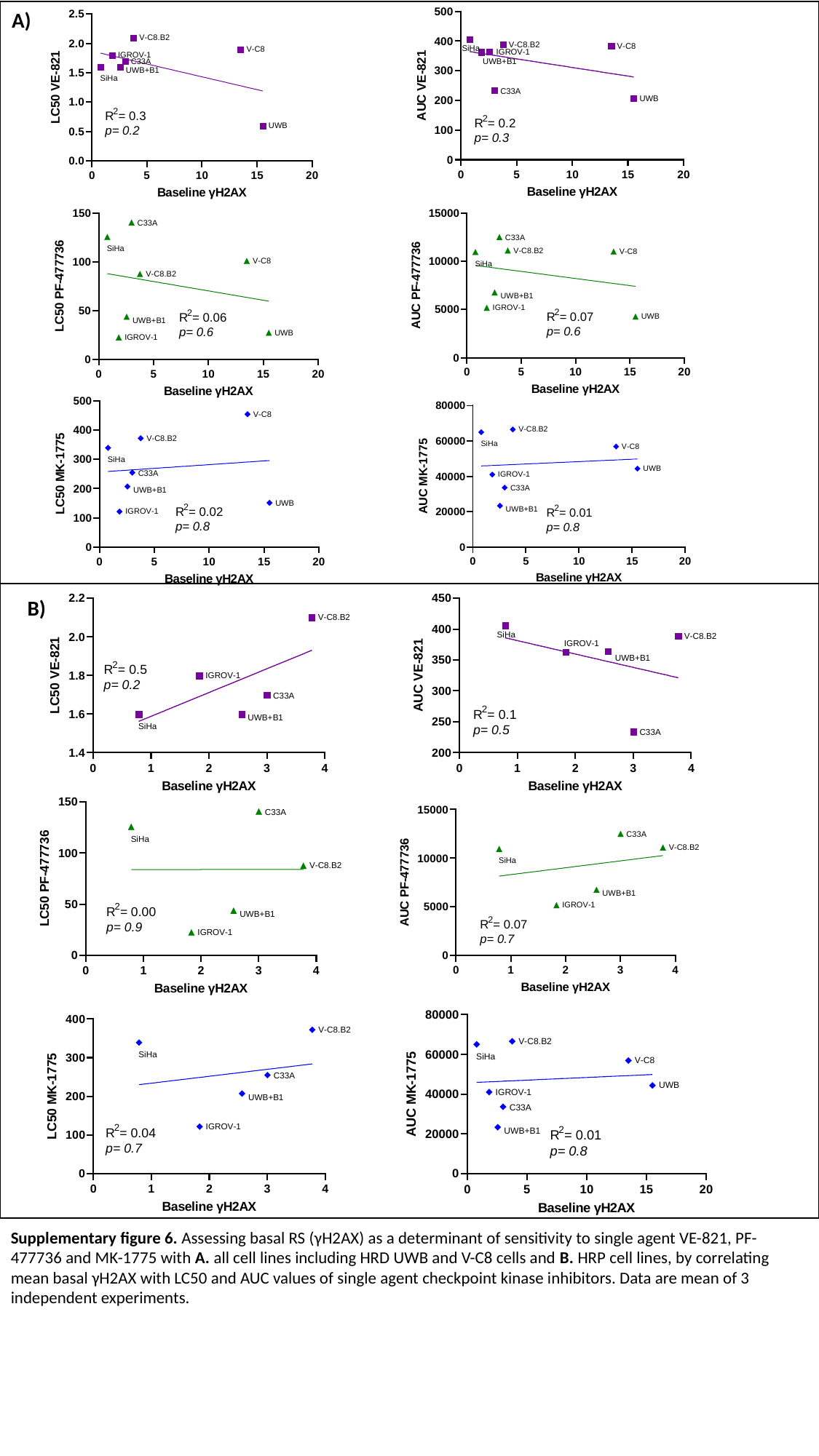

A)
B)
Supplementary figure 6. Assessing basal RS (γH2AX) as a determinant of sensitivity to single agent VE-821, PF-477736 and MK-1775 with A. all cell lines including HRD UWB and V-C8 cells and B. HRP cell lines, by correlating mean basal γH2AX with LC50 and AUC values of single agent checkpoint kinase inhibitors. Data are mean of 3 independent experiments.

### Slide 7
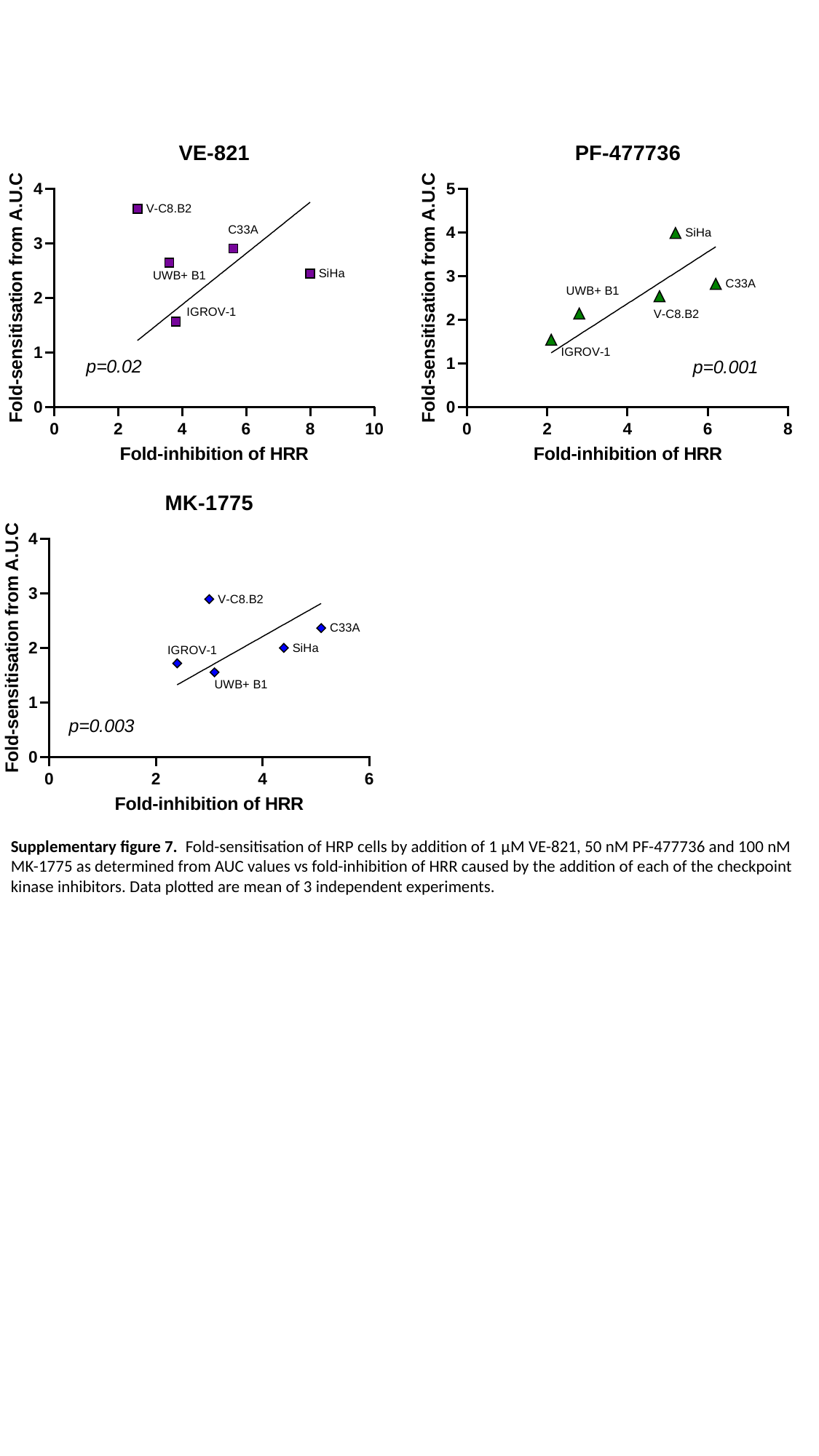

Supplementary figure 7. Fold-sensitisation of HRP cells by addition of 1 µM VE-821, 50 nM PF-477736 and 100 nM MK-1775 as determined from AUC values vs fold-inhibition of HRR caused by the addition of each of the checkpoint kinase inhibitors. Data plotted are mean of 3 independent experiments.
